# Picky – a simple online method designer for targeted proteomics

**DOI:** 10.1101/163154

**Authors:** Henrik Zauber, Marieluise Kirchner, Matthias Selbach

**Affiliations:** Proteome Dynamics, Max Delbrück Center for Molecular Medicine, Robert-Rössle-Str. 10, D-13092 Berlin, Germany; BIH Core Facility Proteomics, Berlin Institute of Health, Anna-Louisa-Karsch-Str. 2, D-10178 Berlin, Germany

## Abstract

Targeted proteomic approaches like selected reaction monitoring (SRM) and parallel reaction monitoring (PRM) are increasingly popular because they enable sensitive and rapid analysis of a preselected set of proteins^1-3^. However, developing targeted assays is tedious and requires the selection, synthesis and mass spectrometric analysis of candidate peptides before the actual samples can be analyzed. The SRMatlas and ProteomeTools projects recently published fragmentation spectra of synthetic peptides covering the entire human proteome^4,5^. These datasets provide very valuable resources. However, extracting the relevant data for selected proteins of interest is not straightforward. For example, developing scheduled acquisition methods (i.e. analyzing specific peptides in defined elution time windows) is complicated and requires adjustments to specific chromatographic conditions employed. Moreover, the number of peptide candidates to be targeted in parallel often exceeds the analytical abilities of the mass spectrometer. In this case, the key question is which peptides can be omitted without losing too much information. Ideally, a method design tool would automatically select the most informative peptides in each retention time window. Until now, none of the available tools automatically generates such optimized scheduled SRM and PRM methods (Figure S1).

---

Here, we present Picky (https://picky.mdc-berlin.de): a fast and easy to use online tool to design scheduled PRM/SRM assays (Figure 1). Users only need to provide identifiers for human (or mouse) proteins of interest. Based on this input, Picky selects corresponding tryptic peptides and their experimentally observed retention times (RTs) from the ProteomeTools dataset for targeted analysis. Picky comes with a scheduling algorithm that adapts to different HPLC gradients (see Figure S2). To this end, users can provide a list of experimentally observed peptide RTs on their HPLC system. A simple shotgun analysis of any standard sample will generate such a list. Picky uses these data to rescale the experimentally observed RTs from ProteomeTools and thus to predict their RTs under the chromatographic conditions employed. About 80% of RTs are correctly predicted within a time window of +/- 3 min (Figure S3), considerably outperforming predictions based on hydrophobicity scores alone (Figure S4). Instead of predicting RTs, users can also directly provide experimentally observed RTs of peptides to be targeted (see Method description). Importantly, the resulting scheduled acquisition list is further optimized if the number of peptides monitored in parallel exceeds a user defined threshold. In this case, the lowest scoring peptide from the protein with the highest number of targeted peptides is removed (Figure S2). This step is repeated until the number of peptides to be targeted in parallel has reached the desired threshold. Hence, Picky selects the best set of peptides covering the proteins of interest under the constraints of the HPLC gradient employed. Parameters such as fragmentation types, charge states, isotope labels and RT window size can be adjusted by the user. For SRM, Picky selects transitions based on the most intense fragment ions observed. Optionally, SRM dwell times can be adjusted according to different protein abundances (Figure S5). The tool exports an inclusion list, which can be imported into the acquisition software of different types of mass spectrometers. In addition, Picky displays and exports annotated fragmentation spectra and a spectral library for all targeted peptides. This library can be imported into Skyline^6^ to validate the acquired SRM/PRM data via intensity correlation methods.

**Fig. 1:**
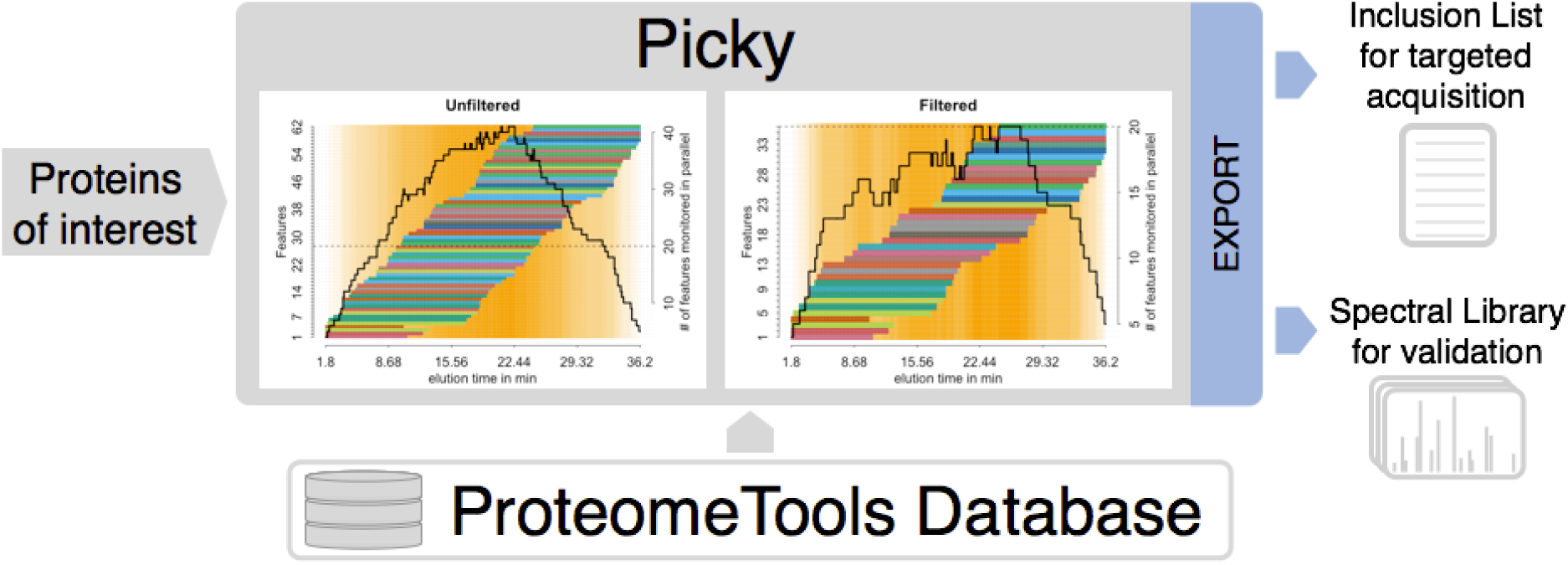
Picky designs targeted acquisition methods (PRM/SRM) for proteins of interest by extracting data from pre-compiled ProteomeTools data. Filtering by the maximal number of co-eluting features selects the best set of peptides for the proteins of interest. Picky exports an inclusion list (for acquisition) and spectral information (for validation) and supports a wide range of mass spectrometers.

To assess the performance of PRM methods designed by Picky we carried out a benchmark experiment. As reference samples we used different amounts of human proteins spiked into 1.4 μg yeast lysate. We only provided Picky with identifiers of proteins to be targeted and a list of experimentally observed peptide retention times. Based on this input, Picky designed an optimized PRM method in less than a minute. We then used this method to analyse the reference samples by PRM. For comparison, we also analyzed the same samples via standard data dependent acquisition (DDA). PRM markedly outperformed DDA at higher dilutions of the spiked-in proteins (Figure S6). For example, at 300 attomoles PRM still identified 31 of the 45 targeted proteins while DDA detected only four. We also targeted the same number of randomly selected human proteins and did not observe a single false-positive hit (Figure S6). Thus, Picky enables detection of human proteins with high sensitivity and specificity.

SRM/PRM data is typically validated by monitoring the chromatographic coelution of multiple transitions for a given peptide^6^. This approach yielded convincing data for high amounts of spiked in proteins but somewhat unclear results for lower amounts (Figure S7). We therefore also compared the PRM data to annotated fragmentation spectra of corresponding synthetic peptides exported by Picky. The high similarity between the spectra (normalized spectrum contrast angle ≥ 0.5) further validated the PRM data (Figure S8). We also compared all acquired UPS1-derived spectra with all fragmentation spectra in the Picky database (Figure S9). We did not observe a single false match with at least five transitions. Hence, Picky enables targeted protein identification with extremely high confidence.

In summary, the Picky tool (i) automatically generates optimized and scheduled SRM/PRM assays for proteins of interest and (ii) provides means to validate the data via known fragmentation spectra of corresponding synthetic peptides. Our benchmark experiment shows that Picky quickly generates an acquisition method that significantly outperforms non-targeted analysis. Picky thus greatly facilitates the targeted analysis of the human (and mouse) proteome.

## Supplemental Information

**Fig. S1:**
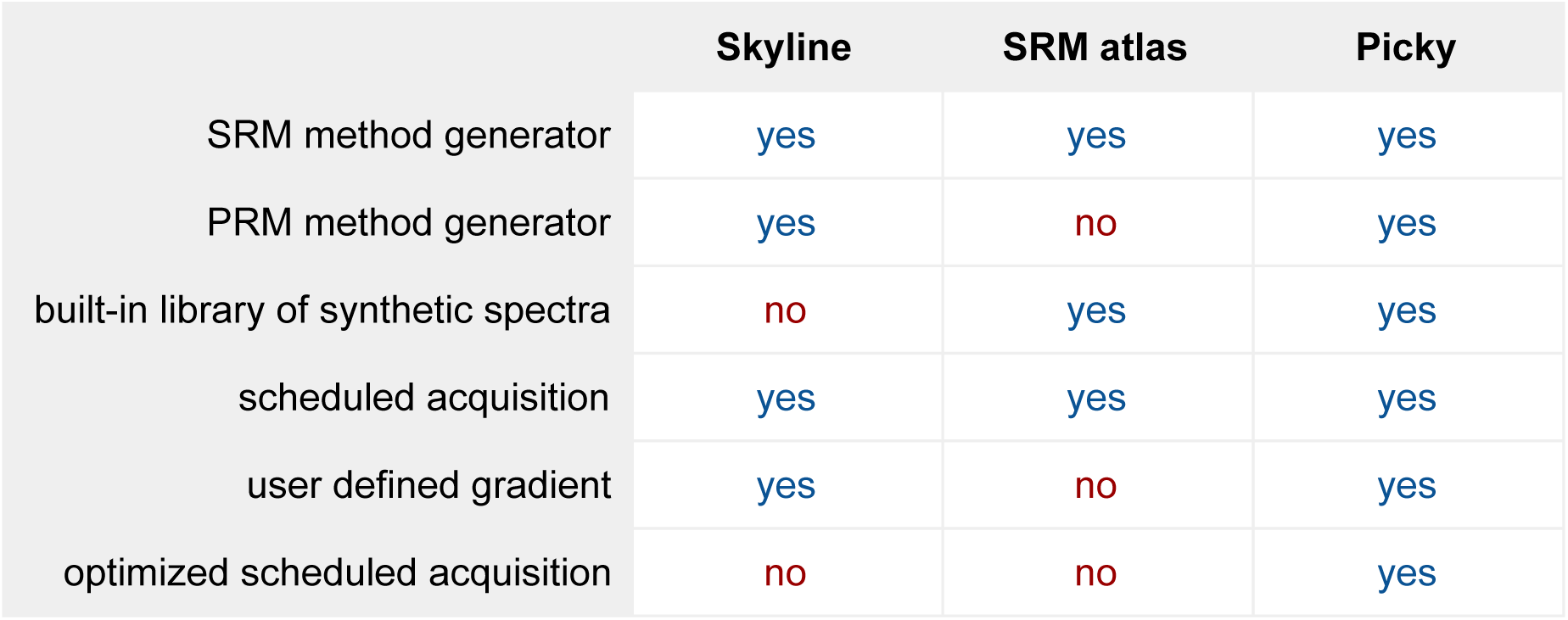
Comparison between different available SRM or PRM method generators.

**Fig. S2:**
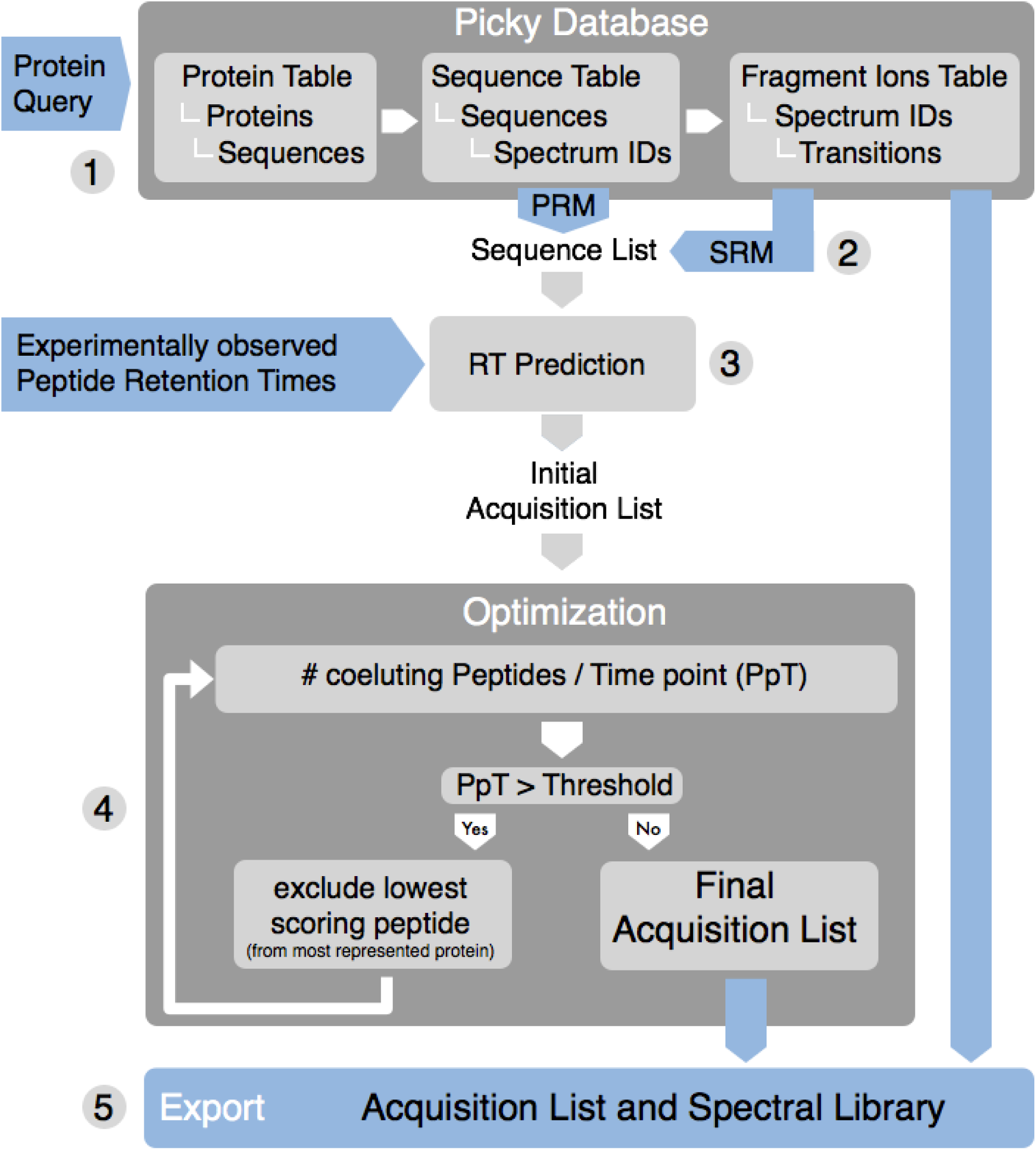
Flowchart of the Picky Algorithm. For more details see section “Picky algorithm” in the Method description.

**Fig. S3:**
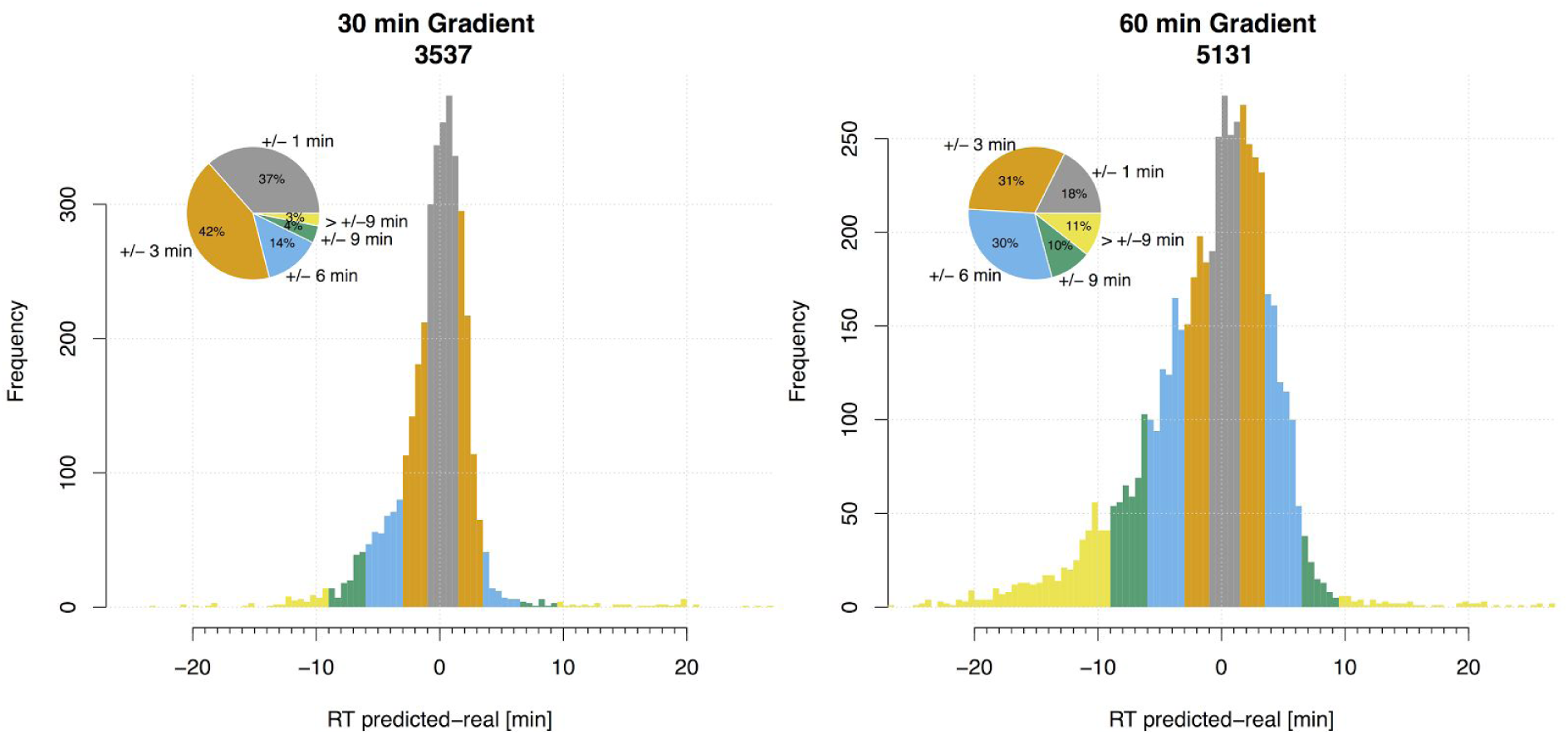
Performance of peptide retention time (RT) prediction implemented in Picky. Differences between observed and predicted RTs based on the rescaled experimentally determined RTs from ProteomeTools. 79% of RTs are correctly predicted within +/- 3 min (or+/- 6) min tolerance in a 30 min (or 60 min) HPLC gradient. The number of peptides analyzed is shown in the title.

**Fig. S4:**
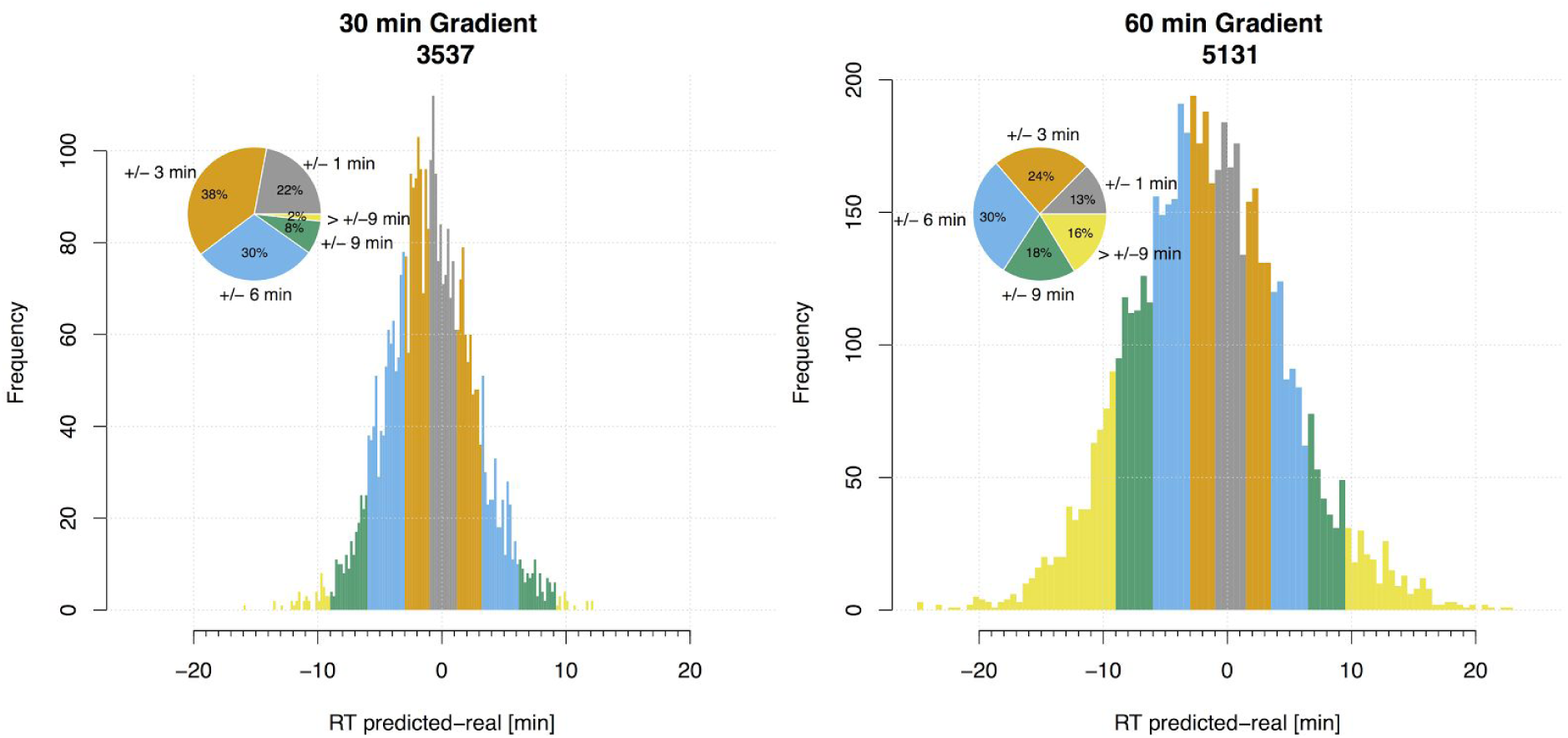
Same as in Fig. S3 but with predicted RTs based on hydrophobicity scores. Predictions based on hydrophobicity scores alone are considerably less accurate than predictions based on experimental RTs (compare to Fig. S3).

**Fig S5:**
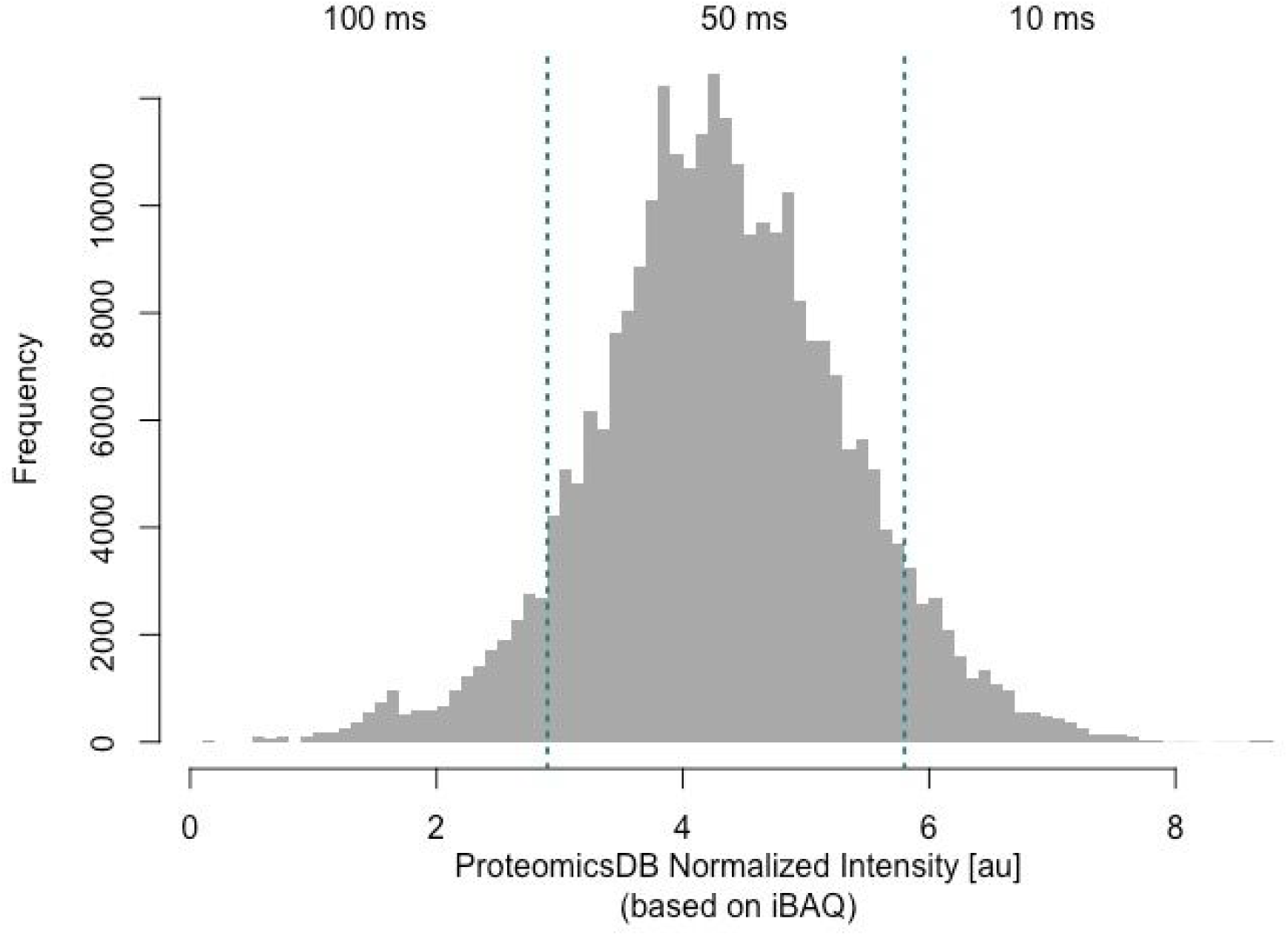
Protein abundance distribution from ProteomicsDB (based on iBAQ values). The abundance range was divided into three bins (divided by turquoise lines) to assign the depicted protein abundance-specific dwell times in Picky. Peptides of proteins not listed in ProteomicsDB receive a dwell time of 100 ms.

**Fig. S6:**
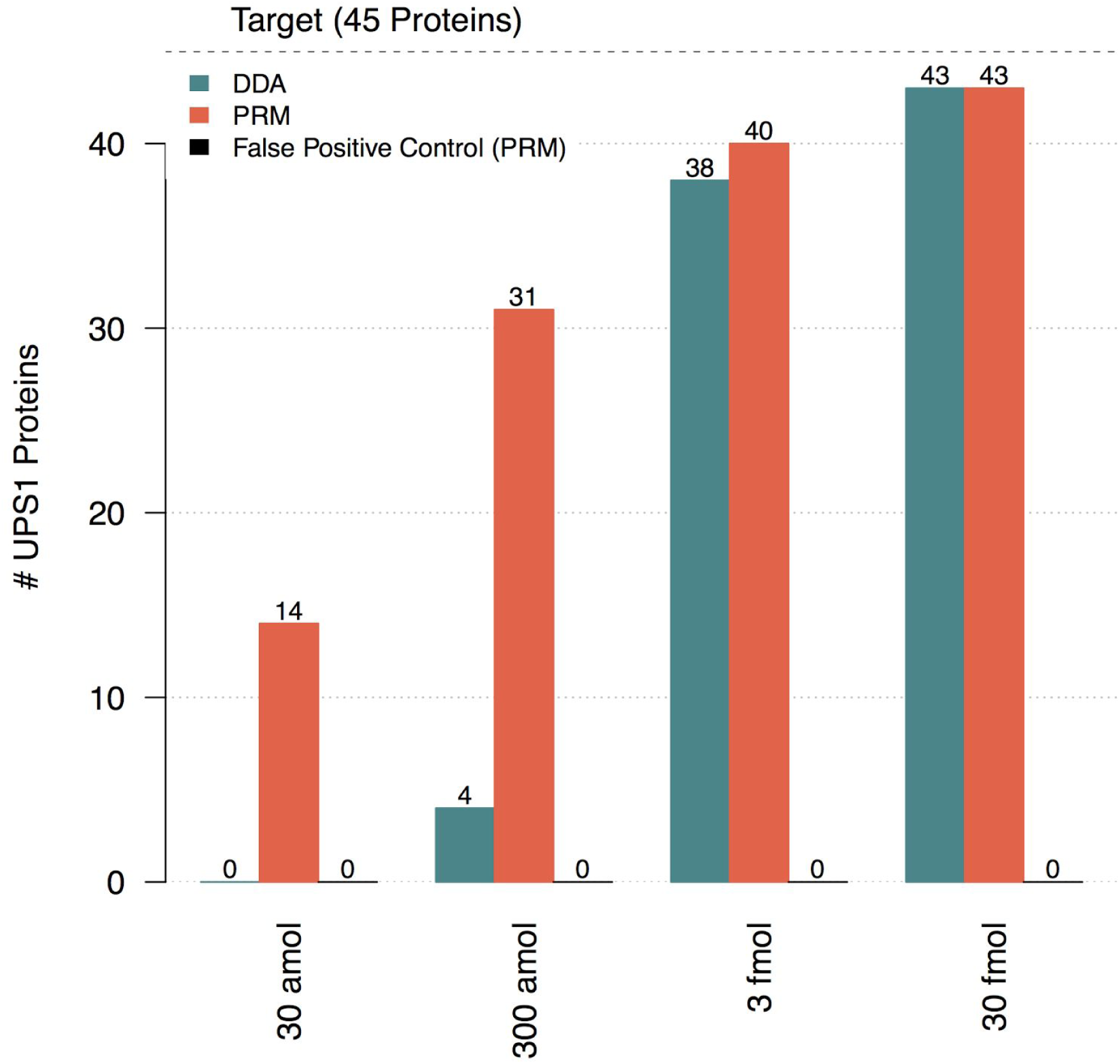
Benchmark experiment to assess the specificity and sensitivity of PRM methods designed by Picky. As a reference sample different amounts of human proteins (UPS1) were spiked into 1.4 μg yeast lysate. A targeted method to detect all human proteins was designed by Picky (see Methods). To control false positives we targeted the same number of randomly selected human proteins (i.e. proteins not actually present in the sample). All samples were analyzed on the same Q Exactive Plus instrument via PRM and DDA. PRM markedly outperformed DDA without giving rise to false positive identifications. Note that we excluded three of the 48 human proteins in UPS1 since they share tryptic peptides with yeast proteins.

**Fig. S7:**
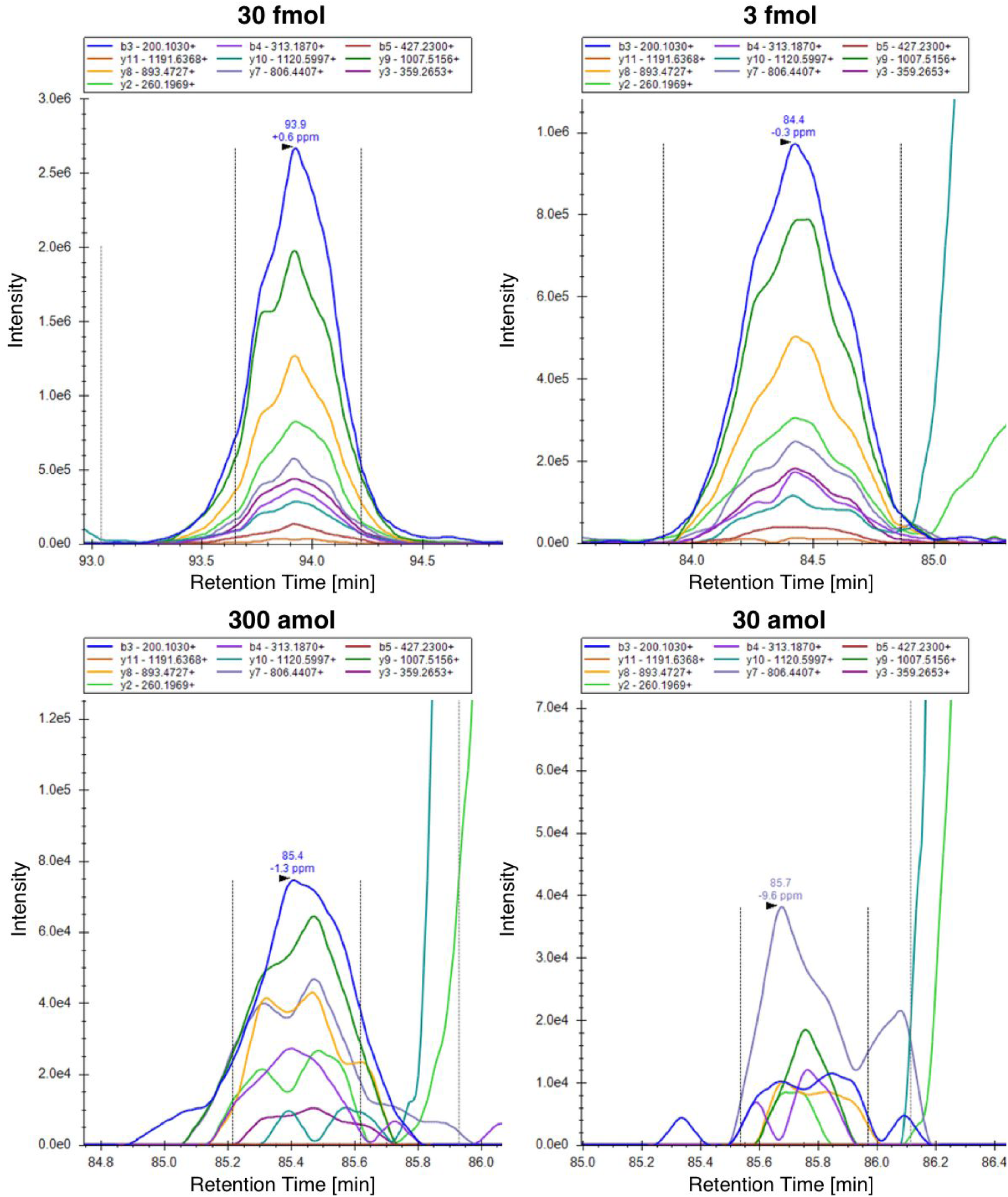
Peaks of the peptide AGALNSNDAFVLK from the protein GSN. Figures were exported from Skyline for the four spike-in amounts 30 fmol, 3 fmol, 300 amol and 30 amol.

**Fig. S8:**
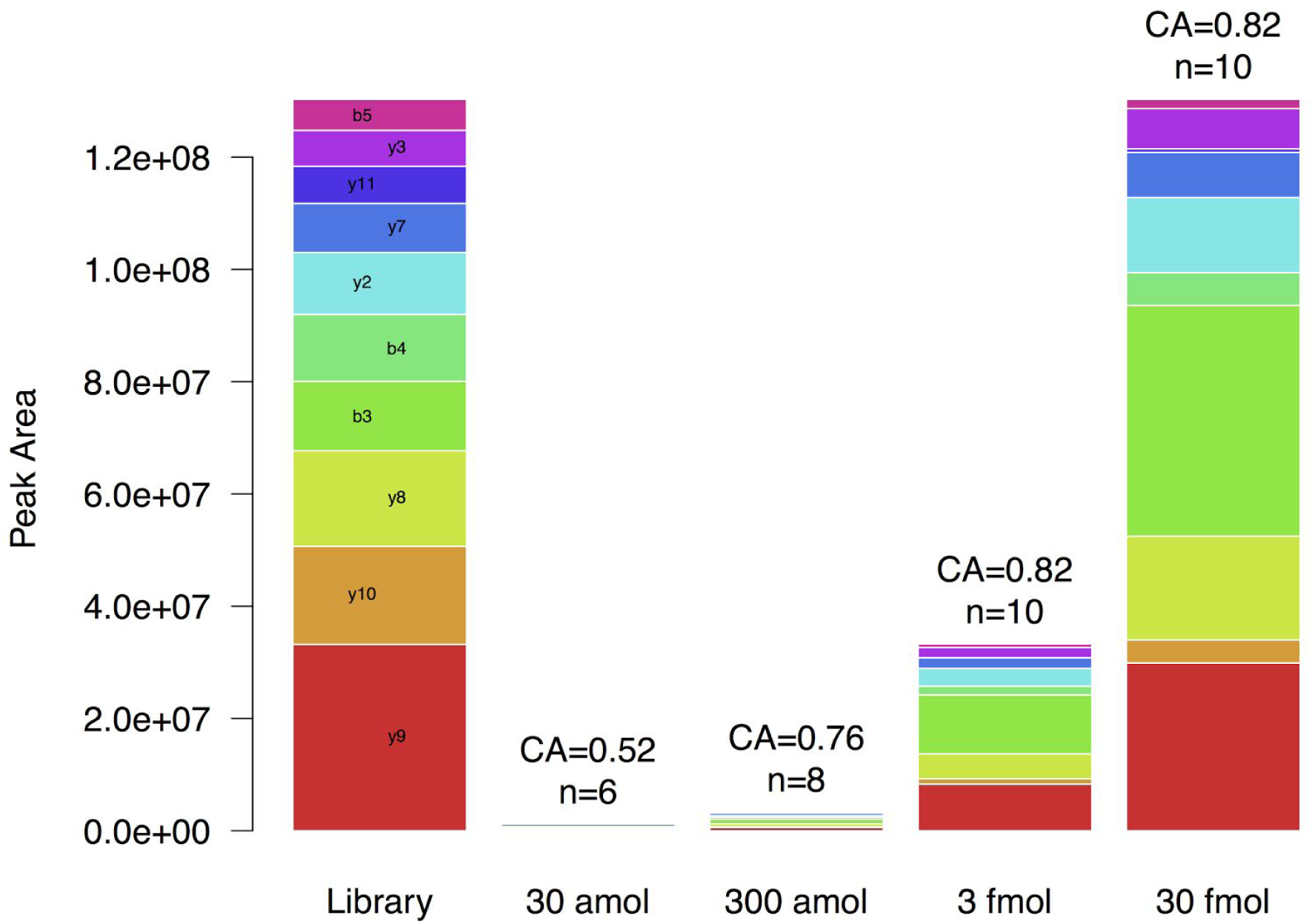
Peak Areas of the peptide AGALNSNDAFVLK from the UPS1 protein GSN at different spike-in amounts (related to Fig. S4). The normalized spectrum contrast angle (CA) and the number of matched transitions is depicted above each stack and indicates spectrum similarity with the library spectrum. The different colors represent the different fragment ions. Library intensities were scaled to the maximal stack sum.

**Fig. S9:**
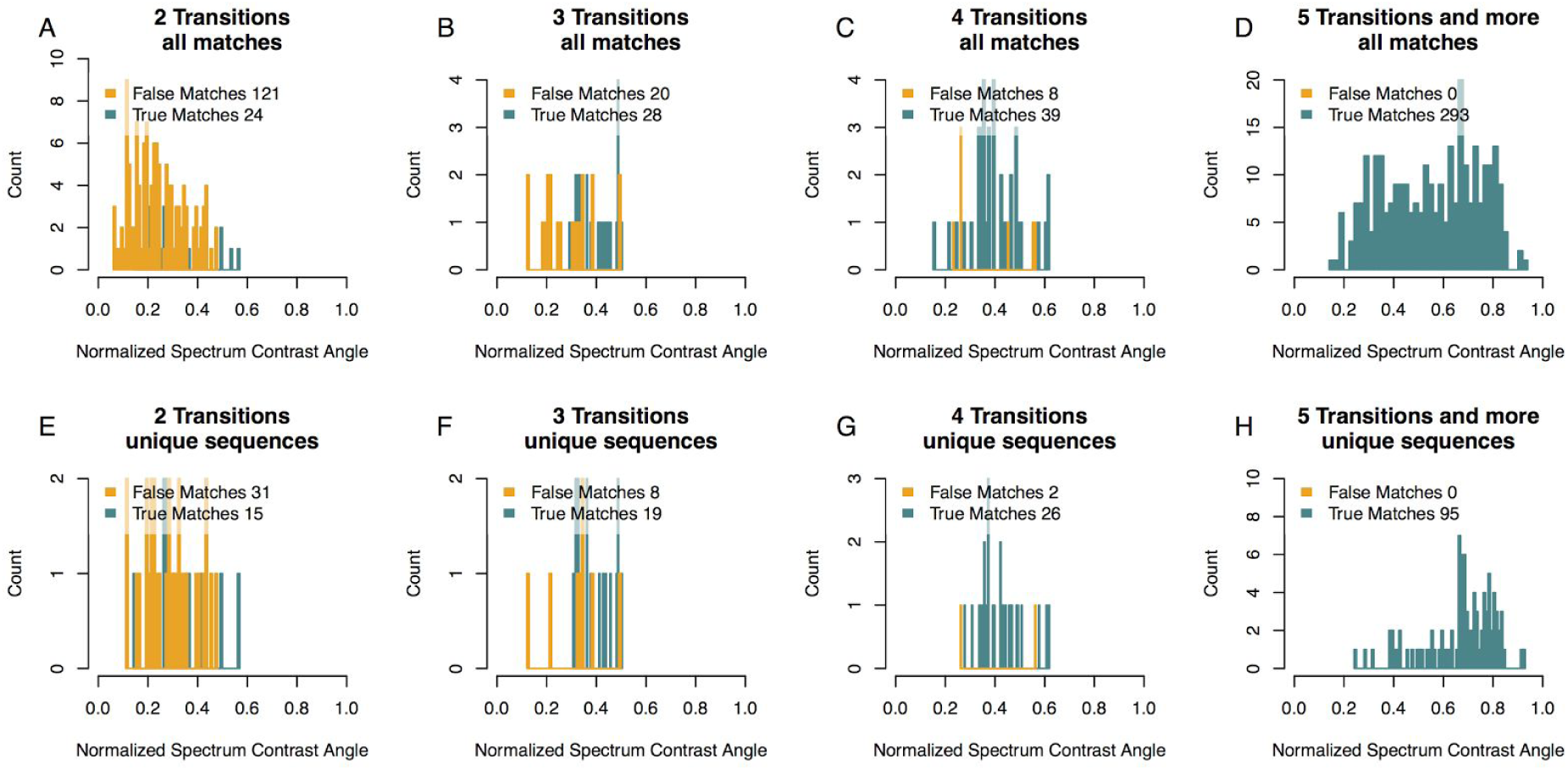
Cross spectrum comparisons between the Picky library and all experimentally observed spectra from peptides of all UPS1 proteins at all concentrations in the benchmark dataset. The normalized spectrum contrast angle (CA) was calculated between spectra with matching precursor and transition masses (20 ppm mass accuracy). True and false matches for different numbers of transitions are shown (turquoise and orange, respectively). With at least five transitions no false match is observed. The top row shows results for all matches (A-D) while the bottom row depicts the highest CA for every unique sequence (E-H).

## Method description

### Picky Database

Data from ProteomeTools was precompiled using msms.txt text files from the available MaxQuant result files. For each peptide species and method-type the best scoring spectrum was picked. We found for almost all proteins listed in ProteomeTools at least one identification event in the provided msms.txt files while only 57 were without any identification event (based on unique gene names). Peptide species and method-types were distinguished by modification, charge, fragmentation type and collision energy. The corresponding raw fragmentation-spectra were extracted from raw-files with a python script using the Thermo MSFileReader and the MSFileReader.py bindings written by François Allen. The data was split into three tables holding information about proteins, peptides and corresponding transitions. All three tables were integrated into a SQLite database in R with the R-package RSQLite. The database is embedded in a shiny environment written in R to enable user friendly access. The R-code for Picky is available on github under the url: https://github.com/rallezumbi/Picky.

### Peptide retention time prediction

Peptide retention times (RTs) can be predicted based on their amino acid sequence by calculating a hydrophobicity score^1^. However, such predictions are not very precise and can deviate from actually observed RTs. Therefore, rather than relying on the hydrophobicity score alone, Picky uses experimentally observed peptides RTs from the ProteomeTools data. These experimentally observed RTs still have to be adjusted to the chromatographic conditions employed by the user. To this end, Picky first calculates hydrophobicity scores^1^ for all peptides in ProteomeTools. A polynomial regression with loess (as is implemented in R) is then used to adjust hydrophobicity scores according to the experimentally observed RT of the corresponding peptides (in ProteomeTools). These calculations are done separately for every raw file in the ProteomeTools dataset. Precomputed adjusted hydrophobicity scores for every peptide are stored in the Picky database. In a second step, Picky uses a list of user defined RTs to predict RTs for the chromatographic system employed. Such a list can be obtained by a shotgun proteomic analysis of any complex sample, ideally immediately before the planned targeted acquisition. Picky uses these data to correlate experimentally observed RTs (in the user defined list) with their calculated hydrophobicities (loess based fit with adjustable parameters on the Picky web interface). Finally, this fit is used to predict RTs of peptides to be targeted via their adjusted hydrophobicities.

To assess the accuracy of these predictions, we analyzed 1μg of HeLa sample (Pierce Hela Digest Standard) with two different gradients (30 and 60; Thermo Fusion Mass Spectrometer; top10 method). Peptides identified in these shotgun runs that match to peptides in the Picky database were used to estimate the accuracy of Picky’s RT prediction algorithm. The list of user defined peptides was obtained from a replicate measurement of the same sample and uploaded to Picky. Using experimentally observed RTs from ProteomeTools improved RT prediction from 60 % to 79 % of analysed peptides falling within a +/- 3 min RT window in a 30 min gradient. Similar improvement was visible for the 60 min gradient (Figure S3 and S4).

Alternatively, instead of predicting RTs based on the ProteomeTools data, users can also provide experimentally observed RTs of peptides to be targeted: Whenever a peptide in the “retention time calibration file” is identical to a peptide to be targeted, Picky uses its experimentally observed RT (from the “retention time calibration file”) rather than its predicted RT. Hence, Picky can be used to define RTs in an iterative manner. First, the tool is used to target a subset of proteins with rather wide RT windows. This reveals the actual RTs of corresponding peptides on the HPLC system employed. Second, the observed RTs from several such subsets are combined and added to the “retention time calibration file”. Picky will then design an acquisition method using the experimentally observed RTs.This allows narrower RT windows and thus increases the number of peptides/proteins that can be targeted in a single run.

### Dwell times

For SRM methods Picky selects dwell times based on protein abundance estimates from ProteomicsDB^2^. To this end, the abundance range (based on iBAQ) was split into three equal windows of low, average and high abundant proteins which were assigned to the dwell times 100, 50 and 10 ms respectively (Figure S5). Proteins not identified in ProteomicsDB are considered to be low abundant and therefore assigned to the 100 ms dwell time fraction. Alternatively, users can set a fixed dwell time that is applied to all peptides in the acquisition list.

### Other Species

All sequences were mapped against the mouse proteome (Uniprot July 2017) using an internal R-script. Subsequently, ∼70 000 human peptides from ProteomeTools shared identity with mouse and have a corresponding spectrum listed in the Picky database. Scientists interested in doing SRM/PRM in mouse samples can restrict Picky to this subset by setting the species button to “mouse”.

### Picky algorithm

Picky first collects all available peptide information for queried proteins considering the initial “Database Query” filters (fragmentation types, detector types, charge states, missed cleavage) and “Additional settings” filter (modified peptides, isoform specificity and proteotypic peptides; Fig S2-1). In Picky, arginine or lysine followed by proline is not considered to be a tryptic cleavage site. Further, all spectra are required to have an Andromeda score higher than 50. In case of SRM the highest intense transitions will be picked based on intensity and the set “Additional settings” filters (number of transitions and number of transitions with a m/z higher than the precursor m/z; Fig S2-2). Scheduling of the acquisition list is initialized by uploading a tab delimited table with a peptide-sequence and retention-time column (“Sequence” and “Retention Time”; Fig S2-3). This file can be obtained from any complex proteomic standard sample. Hydrophobicities of these sequences are calculated and fitted to the retention times using polynomial regression with the loess function as is implemented in R. Subsequently, peptides or transitions from queried proteins can be scheduled by predicting the retention time based on their rescaled hydrophobicity scores (see Rescaled Hydrophobicity Section). The resulting “Initial acquisition List” will be further optimized to fit the filter “maximal number of features monitored in parallel” in an iterative fashion (Fig S2-4): Different peptides in the list are scored according to their posterior error probability (PEP; lower is better) as calculated by MaxQuant and listed in ProteomeTools. Among peptides that coelute and exceed the threshold of “Maximal number of features monitored in parallel”, the lowest scoring peptide from the most represented protein(s) is removed from the acquisition list. It is known, that not all peptides are suitable for quantification even if they are proteotypic^3^. For a reliable quantification it is therefore recommended to choose settings that allow to select for at least two peptides per protein. The Picky algorithm facilitates this selection, by keeping at least two peptides per protein as long as other proteins in the list are represented by more than two available peptides. Importantly, when all proteins are only represented by a single peptide at the given elution time, the Picky algorithm will still exclude the lowest scoring peptide to make sure the maximal number of co-eluting features is not exceeded. In this case, the corresponding protein will be removed from the targeted acquisition method. Picky reports if and which proteins are excluded during the optimization procedure. To prevent this from happening, users can either increase the maximal number of features monitored in parallel, decrease the retention time window (while increasing the risk of missing the peptide) or remove proteins from the query. The final acquisition list can be downloaded in different formats together with the corresponding spectra (Fig S2-5). The MaxQuant deconvoluted spectra and raw spectra are compiled into the MaxQuant msms.txt format. Both types of msms.txt files can be imported into Skyline as a peptide search and used for spectrum comparison.

### Sample Collection, Preparation and Measurements

Universal Protein Standard 1 (UPS1) (Sigma Aldrich) was spiked at different amounts (30 amol, 300 amol, 3 fmol and 30 fmol) into 1.4 μg from total yeast protein extract. Yeast proteins were extracted from *S. cerevisiae* (strain BJ2168). Proteins were digested with trypsin and stage-tipped^4^. Peptides were separated on a reverse phase HPLC system using a self packed column (ReproSil-Pur C18-AQ material; Dr. Maisch, GmbH; 3 h gradient; 5 to 75 % Acetonitrile). Peptides were ionized using an ESI source and analyzed on a Q-Exactive plus (Thermo Fisher). Samples were analyzed with a top10 data-dependent mode acquisition method (DDA) and parallel reaction monitoring method (PRM). Each UPS1 dilution was analyzed once for each analysis mode and concentration (DDA, PRM,PRM-False-Positive-Control) resulting in 12 samples. For DDA settings were briefly: Resolution 70 000 for MS1 (target value: 3,000,000 ions; maximum injection time of 20 ms; dynamic exclusion: 30 s); 17,500 for MS2 (maximum ion collection time of 60 ms with a target of reaching 1,000,000 ions; 2 Da isolation width). MS2 in PRM mode were acquired at a resolution of 17,500, AGC target at 200,000 ions, maximum injection time at 50 ms, isolation window 1.6 m/z). Inclusion lists with 118 peptides were obtained from Picky using default settings to target all 48 UPS1 proteins. The maximal number of features monitored in parallel was set to 60 resulting in a cycle time between 3 and 4 seconds. A false positive control inclusion list was generated with Picky. 48 random human proteins different from the UPS1 set were queried in Picky and analyzed using the described settings. The mass spectrometry proteomics data for the benchmark experiment have been deposited to the ProteomeXchange Consortium via the PRIDE^5^ partner repository with the dataset identifier PXD007039.

Retention time benchmarks were performed by analysing 1 μg of HeLa protein digest standard (Pierce) in DDA mode with a 30 and 60 minute gradient. The setup for the mass spectrometric measurements was as described above but applying shorter gradients (5-75 % Acetonitrile in 30 and 60 minutes). The sample were analyzed on an Orbitrap Fusion Tribrid Mass Spectrometer (Thermo) with the following settings: MS1: Orbitrap Resolution 60000; Scan Range 350-2000, AGC target 4e5, maximum injection time 50 ms. MS2: Top20, Orbitrap HCD, Resolution 15000, AGC target 5e4, maximum injection time 50 ms, dynamic exclusion: 30 s.

### Bioinformatic analyses

DDA runs were analyzed with MaxQuant 1.5.8.0^6^ using default settings (multiplicity=0;Enzyme=Trypsin, including cut after proline; Oxidation (M) and N-terminal Acetylation set as variable modifications; carbamidomethylation (C) as fixed modification; database: uniprot yeast database from october 2014 and ups1 database as provided from Sigma Aldrich; Peptide and Protein FDR set to 0.01). UPS1 Proteins were defined as being identified if a protein-group listed a corresponding UPS1 protein at the first position. PRM data was analyzed with Skyline (3.6.0) with the following settings: Precursor Charges 2 to 7; ion charges 1 to 4; Ion types b and y; up to 6 product ions picked; auto-selection of matching transitions enabled; precursor m/z exclusion window = 2; ion match tolerance = 0.05 m/z; method match tolerance = 0.055 m/z; high selectivity extraction enabled; all matching scans were included; Resolving power of MS2 filtering was set to 17,500 at 400 m/z). A run specific spectral library was imported into Skyline using the peptide search import option. The msms.txt file was imported as downloaded from Picky. Each feature was manually validated in all samples by starting from the highest UPS1 spike in. Peaks needed to be in the range with the observed retention time in the highest concentrated UPS1 sample, have at least four matching transitions and a normalized spectral contrast angle (CA)^7^ higher or equal to 0.5. All b and y ions as selected by Skyline were included into the calculation of the CA. Missing ions in recorded spectra were replaced with zero intensity. The observed median CA was 0.8. Final results were exported as a transition report and compared with the proteinGroups.txt from the DDA analysis using the statistical computing language R. Proteins sharing selected peptides with *S. cerevisiae* or sharing a protein-group in the MaxQuant results were excluded from the analysis. Altogether, 45 UPS1 proteins were included in the final comparison.

## Acknowledgements

We thank Frank Büttner and Christian Sommer for their excellent technical support and the setup of the linux server system. We also like to thank Matthias Ziehm and Daniel Perez-Hernandez for intense testing of the Picky user interface as well as three reviewers for their constructive comments.

## Author contributions

H.Z. developed the tool with input from all authors. H.Z. performed and analyzed all PRM experiments. M.K. performed and analyzed all SRM experiments and performed the RT-benchmark experiments. H.Z and M.S wrote the manuscript.

## References

1. Shi, T. et al. Advances in targeted proteomics and applications to biomedical research. Proteomics 16, 2160–2182 (2016).

2. Peterson, A. C., Russell, J. D., Bailey, D. J., Westphall, M. S. & Coon, J. J. Parallel Reaction Monitoring for High Resolution and High Mass Accuracy Quantitative,Targeted Proteomics. Mol. Cell Proteomics 11, 1475–1488 (2012).

3. Picotti, P. & Aebersold, R. Selected reaction monitoring-based proteomics: workflows,potential, pitfalls and future directions. Nat Meth 9, 555–566 (2012).

4. Kusebauch, U. et al. Human SRMAtlas: A Resource of Targeted Assays to Quantify the Complete Human Proteome. Cell 166, 766–778 (2016).

5. Zolg, D. P., Wilhelm, M., Schnatbaum, K. & Zerweck, J. Building ProteomeTools based on a complete synthetic human proteome. Nature Methods (2017). doi:10.1038/nmeth.4153

6. MacLean, B., Tomazela, D. M. & Shulman, N. Skyline: an open source document editor for creating and analyzing targeted proteomics experiments. Bioinformatics 26, 966–968 (2010).

## References

1. Krokhin, O. V. et al. An improved model for prediction of retention times of tryptic peptides in ion pair reversed-phase HPLC: its application to protein peptide mapping by off-line HPLC-MALDI MS. Molecular & Cellular Proteomics: MCP, 3 (9), 908–919 (2004).

2. Wilhelm, M. et al. Mass-spectrometry-based draft of the human proteome. Nature, 509(7502), 582–587 (2014).

3. Brownridge, P. et al. Global absolute quantification of a proteome: Challenges in the deployment of a QconCAT strategy. Proteomics, 11 (15), 2957–2970 (2011).

4. Rappsilber, J. et al. Stop and go extraction tips for matrix-assisted laser desorption/ionization, nanoelectrospray, and LC/MS sample pretreatment in proteomics. Analytical Chemistry, 75 (3), 663–670 (2003).

5. Vizcaíno, J. A. et al. 2016 update of the PRIDE database and its related tools. Nucleic Acids Research, 44(D1), D447–56 (2016).

6. Cox, J., & Mann, M. MaxQuant enables high peptide identification rates,individualized p.p.b.-range mass accuracies and proteome-wide protein quantification. Nature Biotechnology 26(12), 1367–1372 (2008).

7. Toprak, U. H. et al. Conserved peptide fragmentation as a benchmarking tool for mass spectrometers and a discriminating feature for targeted proteomics. Molecular & Cellular Proteomics, 13(8), 2056–2071 (2014).

